# Monitoring biodiversity in the global change era: The importance of herbaria and genetic diversity

**DOI:** 10.1101/2024.05.01.591834

**Authors:** Melissa Viveiros-Moniz, Ana García-Muñoz, Luis Matias, Mohamed Abdelaziz, A. Jesús Muñoz-Pajares

**Affiliations:** Department of Genetics, University of Granada, Granada, Spain; Área de Biodiversidad y Conservación, Departamento de Biología y Geología, Física y Química Inorgánica, Universidad Rey Juan Carlos, Móstoles, Spain; Departamento de Biología Vegetal y Ecología, Facultad de Biología, Universidad de Sevilla, Sevilla, Spain; Research Unit Modeling Nature, Universidad de Granada, E‐18071 Granada, Spain

## Abstract

Climate change is having far-reaching consequences on all living beings, altering ecosystems, habitats, and biodiversity worldwide. Species distributions are shifting or decreasing, with alpine plant species being particularly threatened. Natural population monitoring allows the assessment of the impact of human-induced global changes. However, traditional monitoring strategies based on individual counts may produce delayed signals of biodiversity loss. These approaches overlook the fact that genetic diversity is the fundamental basis for evolutionary processes, as it enables populations to adapt to environmental changes, including those caused by climate change. Here, we draw attention to the use of genetic diversity in monitoring schemes to anticipate negative trends in biodiversity and propose two fundamental methodologies: genomics and the use of herbarium specimens. Firstly, in contrast to genetic markers conventionally used to quantify genetic diversity, such as microsatellite markers, genomic approaches provide a vast amount of data that does not require previous knowledge of the studied organism, making them suitable for the study of non-model species. Secondly, herbaria worldwide serve as excellent sources of plant material for comparative studies across time with their precise chronologically recorded collection data. The accuracy of genetic diversity estimates increases with sample size, therefore a large number of vouchers is ideally required. However, the availability of specimens from the same species and populations in public herbaria is limited. Different strategies to quantify genetic diversity are proposed depending on the number of specimens available and their geographic distribution. Finally, we illustrate the potential of this approach in the most restrictive scenario, where only a few individuals are available, and there is no conspecific reference genome. Even in this restrictive scenario, there are signs of genetic depauperation in an alpine species with a narrow distribution, but not in a widely distributed congeneric.

---

The rapid shift in global weather patterns is impacting the overall balance of ecosystems, including organisms, communities, distribution, abundance, ecological interactions, and genetic diversity. These changes are endangering the survival of numerous species, leading to severe threat to biodiversity preservation. To mitigate these negative effects, it is necessary to implement appropriate management and conservation efforts. The monitoring of biodiversity plays a crucial role in mitigating the effects of climate change on natural populations, as it helps identifying vulnerable species and making informed decisions to improve ecosystem functioning. An ideal monitoring scheme should detect negative trends early enough to develop effective conservation actions. Biodiversity monitoring involves quantifying changes in the richness of living organisms in a given area over time using various methodologies, such as resampling, long-term monitoring, participatory schemes, or satellite mapping. In most cases, monitorization focuses on the observed number of individuals or species in the studied area (Jetz et al., 2019). However, this approach may result in delayed detection of negative trends, such as when a species has partially or completely disappeared from the study region. By monitoring genetic diversity instead of the number of individuals, negative trends can be detected earlier, as the loss of genetic diversity and particular alleles often precedes the disappearance of populations (Figure 1A). Quantifying genetic diversity and comparing values over time provides information about the extent to which populations or species may be threatened. Assessing changes in the genetic diversity of species during the last decades (or even centuries) is feasible by utilising museum-preserved materials. The vast collection of plant specimens collected across space and time available in worldwide herbaria provides a valuable resource for evaluating the impact of the Anthropocene on the genetic diversity of natural populations.

**Figure 1.**
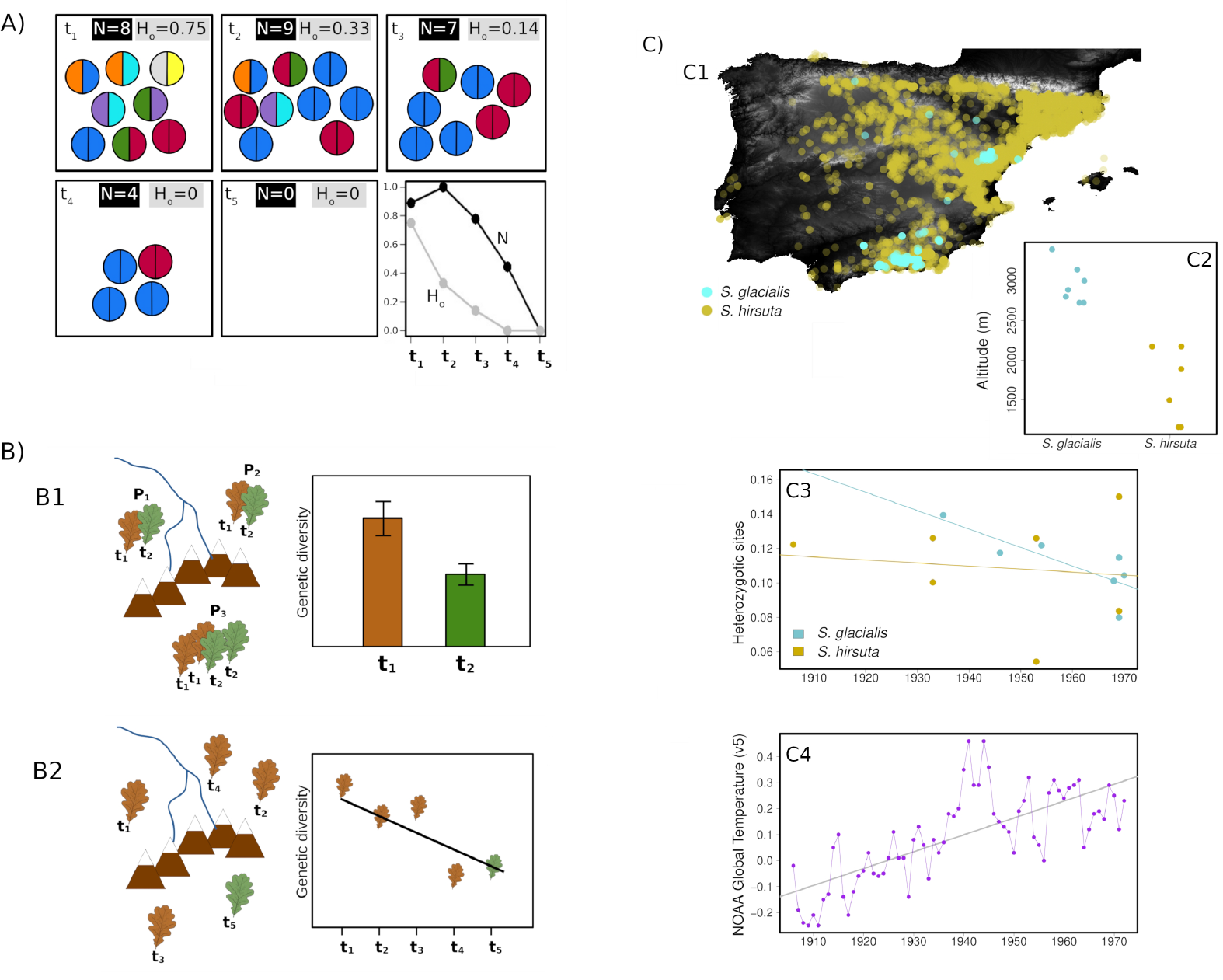
A) Schematic view of the genetic diversity of a theoretical population at five different times (t_1_ to t_5_). Homozygous and heterozygous individuals are represented as circles filled with one and two colours, respectively. For every time, the number of individuals composing the population (N) and the observed heterozygosity (H_o_, estimated as the proportion of heterozygous individuals) are shown. The last plot represents values for N and H_o_ over time. Note the higher sensibility of H_o_ to genetic depauperation B) Alternative strategies proposed to quantify genetic diversity changes over time. B1 compares genetic diversity estimated using a set of herbarium specimens collected at some point in the past (t _1_) with the same number of current individuals (t_2_) collected from the same populations. Statistical support of the differences in genetic diversity is provided by confident intervals estimated using resampling methods. B2 compares individual heterozygosity at different times (t_1_-t_4_ from herbaria and t_5_ from field). Statistical support is provided by a regression analysis. The latter approach is more suitable when the number of herbarium specimens is low to estimate genetic diversity at the population level. C) Temporal genetic diversity variation in two *Sideritis* species. Geographic (C1) and altitudinal (C2) distribution of *Sideritis hirsuta* and *Sideritis glacilalis* in the Iberian Peninsula according to GBIF. C3) Correlation between individual heterozygosity and time for *S. hirsuta* and *S. glacilalis* using samples from the herbarium of the University of Granada collected between 1906 and 1970 (only seven individuals per species available, *S. glacialis* t=-3.00, p-value<0.05; *S. hirsuta* t=-0.28, p-value=0.79). C4) Temperature variation between the same period (1906–1970, data source https://www.eea.europa.eu/data-and-maps/daviz/global-average-air-temperature-anomalies-6/).

## Using genomics and herbarium specimens to detect genetic diversity loss

Herbarium records have become increasingly important in biodiversity research and conservation biology to reliably evaluate changes in species distribution, phenotypic traits, and phenological characteristics driven by climate alterations (Davis et al., 2015, Greve et al., 2016). The potential of applying molecular genetics and genomics tools on herbarium specimens also represent an excellent opportunity to study evolutionary changes directly on the studied populations (Wandeler et al., 2007). In fact, several studies have addressed this question using microsatellite markers. For instance, Gavrilenko et al. (2023) analysed the genetic diversity of Chilean cultivated potato herbarium specimens and present-day gene bank accessions, and the study by Rosche et al. (2022) explored the use of herbarium specimens to track population genetic signatures linked to the local extinction events of *Biscutella laevigata* subsp. *gracilis*. However, there are very fewer studies using whole genome sequencing to quantify genetic diversity using herbarium samples. Nygaard et al. (2022) have recently demonstrated the successful use of SNP genotyping and environmental niche modelling of historical herbarium specimens and modern samples to monitor the genetic diversity of *Dracocephalum ruyschiana* (Lamiaceae).

To examine trends in genetic variation over time, a simple strategy is to quantify genetic diversity using herbarium specimens collected at known locations (P_1_, P_2_, …, P_n_) during specific time in the past (t_1_). This diversity can then be compared to the values obtained from the same number of individuals collected at the same locations during a different time (t_2_). Resampling individual pairs from the genomic dataset using techniques such as bootstrapping or jackknife may be useful to obtain confidence intervals and statistical support for the observed differences in genetic diversity between t_1_ and t_2_ (Figure 1B1). Because it is typically difficult to have enough herbarium individuals collected in a single year to follow this approach, a different strategy can be employed by using all available herbarium specimens of a given species, collected in different times (t_1_, t_2_, …, t_n_) and known locations (P_1_, P_2_, …, P_n_). Trends in genetic diversity can be obtained by plotting intraindividual genetic diversity estimates (heterozygosity) against collection time, and a regression analysis can provide statistical significance values (Figure 1B2, see Supplementary Material for a pipeline in R).

To monitoring genetic diversity, it is often necessary to first characterize the study system. This can be achieved through the design and testing of microsatellite primers (e.g. Gavrilenko et al., 2023; Rosche et al., 2022) or the development of a panel of SNP markers (e.g. Nygaard et al., 2022). Whole genome sequencing approaches may reduce the need for prior knowledge when monitoring poorly characterized species. *De novo* assembly of sequenced reads can easily reconstruct the whole genome sequence of organelles, but identifying variants in the nuclear genome of non-model species may be challenging. In such cases, it is more suitable to align sequenced reads to a reference genome. Although the genetic distance between samples and the reference genome can affect mapping efficiency and genotyping accuracy (Thorburn et al., 2023), this effect is likely to be negligible when the objective is to compare genetic diversity over time because the bias effect is consistent across the samples being compared. Consequently, monitoring nuclear genetic diversity of poorly characterized species can be performed using whole genome sequencing and the reference genome from a relative. The following section oulines the monitoring of two *Sideritis* species lacking a conspecific reference genome available in GenBank and have only a limited number of herbarium-preserved individuals.

## Monitoring the genetic diversity loss of two species of *Sideritis* in Sierra Nevada

The distribution of plant species is the result of complex interactions between multiple factors that typically lead to a fit between environmental conditions and species requirements. In response to rising temperatures, plant species migrate to higher elevations to colonise new areas with more favourable conditions. However, the range of many alpine species is threatened due to habitat loss and the lack of new available areas as they reach mountain summits. The Sierra Nevada mountain range is located in southeastern Spain and is considered to be a plant diversity hotspot in the Mediterranean region. It hosts 33.2% of the total vascular plants found in the Iberian Peninsula, including 98 endemic taxa, 70% of which are threatened by global warming (Benito et al., 2011; Lorite, 2016). Most of the endemisms in Sierra Nevada inhabit the summit areas (Benito et al., 2011), highlighting the importance of monitoring genetic diversity of these species to identify if conservation measures are necessary.

The genus *Sideritis* (Lamiaceae) comprises approximately 140 species, with over one quarter of them found in the Iberian Peninsula. Our monitoring strategy focuses on two *Sideritis* species differing in their distribution ranges. *Sideritis hirsuta* L. has a wide distribution across the Iberian Peninsula, while *S. glacilalis* Boiss is endemic to high-altitude areas in the southeast of the Iberian Peninsula (Figure 1C1). The survey was restricted to specimens collected before 1971 from various populations in Sierra Nevada and preserved in the Herbarium of the University of Granada with known collection date, location, and altitude. DNA was extracted from a total of seven herbarium samples per species. Individual libraries were prepared and sequenced using the Illumina technology and the resulting reads were aligned against the genome of *Ballota nigra* (OX344720.1). We filtered the resulting VCF to retain SNPs with quality values over 20, DP lower than 100 and less than 85% of missing data. Finally, we explored the proportion of heterozygous sites as a metric for intra-individual genetic diversity.

As shown in Figure 1C3 and regardless the low sample size, we found a significant decrease in genetic diversity in *S. glacialis* samples (t=-3.00, p-value<0.05), while *S. hirsuta* has remained stable throughout the studied period (t=-0.28, p-value=0.79). The combination of limited distribution and greater altitudinal range (F=39.35, p-value<0.0001) probably lead to the observed patterns solely in *S. glacialis*. These findings highlight the vulnerability of isolated high-altitude plant communities to genetic diversity loss and emphasize the importance of herbaria as genetic material sources from past populations and whole genome sequencing as a tool (requiring no previous information on the studied system and then suitable for non-model species) to predict the impact of the Anthropocene on biodiversity. The strategies described may aid in comprehending the impcat of climate change on plant species in different environments at a global scale. These methodologies can assist in developing proactive strategies to safeguard the genetic diversity of plant species, thereby preventing the loss of species diversity.

