## Supplementary material for "Monitoring biodiversity in the global change era: The importance of herbaria and genetic diversity": SupplementaryMaterial.html

### 1 Processing fastq files

#### 1.1 Mapping sequences against reference genome

```
# Mapping reads
bwa mem -t 15 reference.fasta Individual1_1.fq.gz Individual1_2.fq.gz > Individual1.sam

# Converting sam file into bam file
samtools sort -o Individual1.bam -O bam -@15 Individual1.sam

# Variant calling and labeling low quality variants
bcftools mpileup -Ou -f reference.fasta Individual1.sam | bcftools call -Ou -mv | bcftools filter -s LowQual -e '%QUAL<20 || DP>100' > Individual1.flt.vcf
```

This was repeated for every individual and the resulting vcf files
were merged into a single vcf as follows:

```
bcftools merge Individual1.flt.vcf Individual2.flt.vcf .... IndividualN.filt.vcf > Allid.vcf
```

#### 1.2 Filtering VCF

```
# Removing low quality variant, indels and missing data
vcftools --vcf Allid.vcf --remove-filtered-all --remove-indels --max-missing 0.85 --recode out Allid_filtered.vcf
```

### 2 Strategy A: Confident intervals using multiple individuals per year

Two filtered vcf must be produced for this strategy: PAST.vcf and
PRESENT.vcf, containing variants for individuals sampled in the past
(time 1 or t1) and variants for individuals sampled in the present (time
2 or t2). For this example, we have produced two vcf files mapping
*Arabidopsis thaliana* reads donwloaded from GenBank (five
individuals per file). The first vcf file (PAST) contains variants
identified in five individuals (accessions SRR1945855, SRR1945917,
SRR1945937, SRR1945986, and SRR1946065) whereas the second file contains
variants identified in two individuals, repeated multiple times to
account for a total of five individuals (accessions SRR1945584 repeated
three times and SRR1945650 repeated twice).

#### 2.1 Estimating genetic diversity

This will produce nucleotide divergency per site across every site in
the two vcf files:

```
vcftools --vcf PAST.vcf --site-pi --out PAST.txt
vcftools --vcf PRESENT.vcf --site-pi --out PRESENT.txt
```

#### 2.2 Estimating confident intervals using jakknife

This produces vcf files with N-1 individual…

..for the past dataset…

```
bcftools view -s id9,id12,id16,id19 -o PASTno7.vcf -O v PAST.vcf
bcftools view -s id7,id12,id16,id19 -o PASTno9.vcf -O v PAST.vcf
bcftools view -s id7,id9,id16,id19 -o PASTno12.vcf -O v PAST.vcf
bcftools view -s id7,id9,id12,id19 -o PASTno16.vcf -O v PAST.vcf
bcftools view -s id7,id9,id12,id16 -o PASTno19.vcf -O v PAST.vcf
```

…and or the present dataset

```
bcftools view -s id1.2,id1.3,id2,id2.2 -o PRESENTno1.vcf -O v PRESENT.vcf
bcftools view -s id1,id1.3,id2,id2.2 -o PRESENTno1.2.vcf -O v PRESENT.vcf
bcftools view -s id1,id1.2,id2,id2.2 -o PRESENTno1.3.vcf -O v PRESENT.vcf
bcftools view -s id1,id1.2,id1.3,id2.2 -o PRESENTno2.vcf -O v PRESENT.vcf
bcftools view -s id1,id1.2,id1.3,id2 -o PRESENTno2.2.vcf -O v PRESENT.vcf
```

This will produce nucleotide divergency per site across every site in
the previous vcf files…

…for the past…

```
vcftools --vcf PASTno7.vcf --site-pi --out PASTno7.txt
vcftools --vcf PASTno9.vcf --site-pi --out PASTno9.txt
vcftools --vcf PASTno12.vcf --site-pi --out PASTno12.txt
vcftools --vcf PASTno16.vcf --site-pi --out PASTno16.txt
vcftools --vcf PASTno19.vcf --site-pi --out PASTno19.txt
```

…and for the present

```
vcftools --vcf PRESENTno1.vcf --site-pi --out PRESENTno1.txt
vcftools --vcf PRESENTno1.2.vcf --site-pi --out PRESENTno1.2.txt
vcftools --vcf PRESENTno1.3.vcf --site-pi --out PRESENTno1.3.txt
vcftools --vcf PRESENTno2.vcf --site-pi --out PRESENTno2.txt
vcftools --vcf PRESENTno2.2.vcf --site-pi --out PRESENTno2.1.txt
```

#### 2.3 Graphs

```
 # Reading estimated diversity values per site and computing the average...

 #... for the past...
PI_PAST<-read.table(file="PAST.txt.sites.pi",sep="\t",header=T)
REAL<-mean(PI_PAST[,3],na.rm=T)

#... and for the present.
PI_PRESENT<-read.table(file="PRESENT.txt.sites.pi",sep="\t",header=T)
REAL<-c(REAL,mean(PI_PRESENT[,3],na.rm=T))

# Reading jackknife...

#...for the past...
PI_PASTno7<-read.table(file="PASTno7.txt.sites.pi",sep="\t",header=T)
JACKpast<-mean(PI_PASTno7[,3],na.rm=T)
PI_PASTno9<-read.table(file="PASTno9.txt.sites.pi",sep="\t",header=T)
JACKpast<-c(JACKpast,mean(PI_PASTno9[,3],na.rm=T))
PI_PASTno12<-read.table(file="PASTno12.txt.sites.pi",sep="\t",header=T)
JACKpast<-c(JACKpast,mean(PI_PASTno12[,3],na.rm=T))
PI_PASTno16<-read.table(file="PASTno16.txt.sites.pi",sep="\t",header=T)
JACKpast<-c(JACKpast,mean(PI_PASTno16[,3],na.rm=T))
PI_PASTno19<-read.table(file="PASTno19.txt.sites.pi",sep="\t",header=T)
JACKpast<-c(JACKpast,mean(PI_PASTno19[,3],na.rm=T))

#... and for the present.
PI_PRESENTno1<-read.table(file="PRESENTno1.txt.sites.pi",sep="\t",header=T)
JACKpresent<-mean(PI_PRESENTno1[,3],na.rm=T)
PI_PRESENTno1.2<-read.table(file="PRESENTno1.2.txt.sites.pi",sep="\t",header=T)
JACKpresent<-c(JACKpresent,mean(PI_PRESENTno1.2[,3],na.rm=T))
PI_PRESENTno1.3<-read.table(file="PRESENTno1.3.txt.sites.pi",sep="\t",header=T)
JACKpresent<-c(JACKpresent,mean(PI_PRESENTno1.3[,3],na.rm=T))
PI_PRESENTno2<-read.table(file="PRESENTno2.txt.sites.pi",sep="\t",header=T)
JACKpresent<-c(JACKpresent,mean(PI_PRESENTno2[,3],na.rm=T))
PI_PRESENTno2.1<-read.table(file="PRESENTno2.1.txt.sites.pi",sep="\t",header=T)
JACKpresent<-c(JACKpresent,mean(PI_PRESENTno2.1[,3],na.rm=T))

# Estimating confident intervals as 1.96 times the ration between standard deviation and the square root of sample size
Interval_present <- 1.96*sd(JACKpresent)/sqrt(length(JACKpresent))
Interval_past <- 1.96*sd(JACKpast)/sqrt(length(JACKpast))

# Plotting bars and confident intervals
vals<-barplot(REAL,ylim=c(0,0.2),col=c("#b46517","#468a1a"),las=2)
arrows(x0=vals[1],x1=vals[1],y0=REAL[1] - Interval_past,y1=REAL[1] + Interval_past,angle=90,code=3)
arrows(x0=vals[2],x1=vals[2],y0=REAL[2] - Interval_present,y1=REAL[2] + Interval_present,angle=90,code=3)
axis(1,at=vals,labels=c("t1","t2"),cex=1.7)
```

### 3 Strategy B: Regression using few individuals per year

#### 3.1 Estimating intraindividual genetic diversity

This produces a summary for each individual of the observed number of
homozygous sites, the expected number of homozygous sites, the total
number of sites that the individual has data for and the inbreeding
coefficient F.

```
vcftools --vcf Allid_filtered.vcf --het --out sideritis.txt
```

#### 3.2 Graphs

```
# This reads the output of the vcftools --het command above.
side<-read.table("sideritis.txt",sep="",header=T)

# Providing the proper name to every sample...
Species<-c("glacialis","glacialis","glacialis","glacialis","glacialis","glacialis","glacialis","hirsuta","hirsuta","hirsuta","hirsuta","hirsuta","hirsuta","hirsuta")
side$Species<-Species

# ...and also the year and elevation (if available)
side$SpeciesYear<-c(1969,1969,1970,1954,1935,1946,1968,1969,1953,1969,1953,1933,1933,1906)
side$Altitude<-c(2724,2724,2800,3141,2887,3000,3396,NA,2173,1888,2173,1156,1156,1495)


# This estimates heterozygosity using homozygosity values and the total number of sites
side$het <- (side$N_SITES - side$O.HOM.)/side$N_SITES
side$het.expected <- (side$N_SITES - side$E.HOM.)/side$N_SITES

#Genetic diversity across time

side$Species<-as.factor(side$Species)
plot(side$Year,side$het,xlab="Year",cex=2,ylab="Proportion of observed heterozygous sites",pch=16,main="Genetic diversity across time",col=c("#7AC5CD","gold3")[as.numeric(side$Species)])
legend(x = "bottomleft", legend = c(expression(italic("S. glacialis")),expression(italic("S. hirsuta"))), fill = c("#7AC5CD","gold3"),bty="n")

glaci<-side[which(side$Species=="glacialis"),]
cor.test(glaci$Year,glaci$het)
```

```
## 
##  Pearson's product-moment correlation
## 
## data:  glaci$Year and glaci$het
## t = -3.0079, df = 5, p-value = 0.02983
## alternative hypothesis: true correlation is not equal to 0
## 95 percent confidence interval:
##  -0.9696068 -0.1250637
## sample estimates:
##        cor 
## -0.8025385
```

```
abline(lm(glaci$het~glaci$Year),col="#7AC5CD",lwd=2)

hirsuta<-side[which(side$Species=="hirsuta"),]
cor.test(hirsuta$Year,hirsuta$het)
```

```
## 
##  Pearson's product-moment correlation
## 
## data:  hirsuta$Year and hirsuta$het
## t = -0.28047, df = 5, p-value = 0.7904
## alternative hypothesis: true correlation is not equal to 0
## 95 percent confidence interval:
##  -0.8023181  0.6936099
## sample estimates:
##        cor 
## -0.1244542
```

```
abline(lm(hirsuta$het~hirsuta$Year),col="gold3",lwd=2)
```

```
#Elevation range plot

stripchart(side$Altitude ~ side$Species, group.names = c("","") ,cex.main=1.5,cex.lab=1.2,pch = 19, method = "jitter",cex=2, col.lab="#6C4B5E",
           jitter = 0.2, main="Elevation range",xlab="Species",ylab="Elevation (m)", vertical = TRUE, col =c("#7AC5CD","gold3"))
#text(2.157794,3215.335,substitute(paste(italic("n.s."))))

axis(1, font.axis=3,at=1:2,cex=3,labels = c("S. glacialis","S. hirsuta") )
```
